# Statistical inference of mechanistic models from qualitative data using an efficient optimal scaling approach

**DOI:** 10.1101/848648

**Authors:** Leonard Schmiester, Daniel Weindl, Jan Hasenauer

## Abstract

Quantitative dynamical models facilitate the understanding of biological processes and the prediction of their dynamics. These models usually comprise unknown parameters, which have to be inferred from experimental data. For quantitative experimental data, there are several methods and software tools available. However, for qualitative data the available approaches are limited and computationally demanding.

Here, we consider the optimal scaling method which has been developed in statistics for categorical data and has been applied to dynamical systems. This approach turns qualitative variables into quantitative ones, accounting for constraints on their relation. We derive a reduced formulation for the optimization problem defining the optimal scaling. The reduced formulation possesses the same optimal points as the established formulation but requires less degrees of freedom. Parameter estimation for dynamical models of cellular pathways revealed that the reduced formulation improves the robustness and convergence of optimizers. This resulted in substantially reduced computation times.

We implemented the proposed approach in the open-source Python Parameter EStimation TOolbox (pyPESTO) to facilitate reuse and extension. The proposed approach enables efficient parameterization of quantitative dynamical models using qualitative data.

## 1 Introduction

In systems and computational biology, quantitative dynamical models based on ordinary differential equations (ODEs) are widely used to study cellular processes (Klipp *et al.*, 2005; Aldridge *et al.*, 2006; Schöberl *et al.*, 2009; Bachmann *et al.*, 2011). Unknown parameters of these ODE models are often inferred from experimental data (Banga, 2008; Raue *et al.*, 2013a). This is done by minimizing the distance between measured data and model simulation, e.g. the mean squared error, the mean absolute error or the maximum likelihood (Raue *et al.*, 2013a). However, not all experimental techniques and setups provide quantitative data that allow for a direct comparison of measured and simulated data.

In many experimental setups, the measured values only provide information about the qualitative behaviour, e.g. that some quantity decreases or increases. Frequently encountered reasons are (i) unknown nonlinear dependencies of the measured signal on the internal state of the system, e.g. for Förster resonance energy transfer (FRET) (Birtwistle *et al.*, 2011) and (ii) detection thresholds and saturation effects, e.g. for Western blotting (if not properly designed) (Butler *et al.*, 2019). For these techniques a specific fold change in the measured signal does not imply the same fold change in the measured species. Yet, there is a monotonic relation between measured species and signal, meaning that – if the measurement noise is neglected – the ordering is still preserved.

The use of qualitative data is not supported by established parameter estimation toolboxes such as AMIGO (Balsa-Canto & Banga, 2011), COPASI (Hoops *et al.*, 2006), Data2Dynamics (Raue *et al.*, 2015), and PESTO (Stapor *et al.*, 2018) (along with its Python reimplementation pyPESTO (Schälte *et al.*, 2019)). However, two methods have been proposed which facilitate the use of qualitative data in dynamical systems: (1) Mitra *et al.* (2018) used an *ad hoc* approach based on the formulation of qualitative data as inequality constraints. The degree to which the inequality constraints were violated was used as objective function. The parameters were estimated by minimizing this penalized objective function. This approach was implemented in the toolbox py-BioNetFit (Mitra *et al.*, 2019) and recently extended using a probabilistic distance measure (Mitra & Hlavacek, 2019). (2) Pargett & Umulis (2013) and Pargett *et al.* (2014) used the concept of *optimal scaling* established in statistics (Shepard, 1962). Instead of imposing inequality constraints, the optimal scaling method determines the best quantitative representation of the qualitative data. This quantitative representation is referred to as *surrogate data.* The parameters are estimated by splitting the optimization of the parameters in an outer and an inner problem (Figure 1A). In the outer problem the model parameters of the dynamical model are optimized given the parameter-dependent optimal surrogate data computed in the inner problem by minimizing the difference between surrogate data and model simulation. In the inner optimization, the optimal surrogate data for a given model simulation are determined, such that inconsistencies of the model simulation with the qualitative measurement data are penalized. While the optimal scaling approach is deeply grounded in statistical theory, it is computationally demanding.

**Figure 1:**
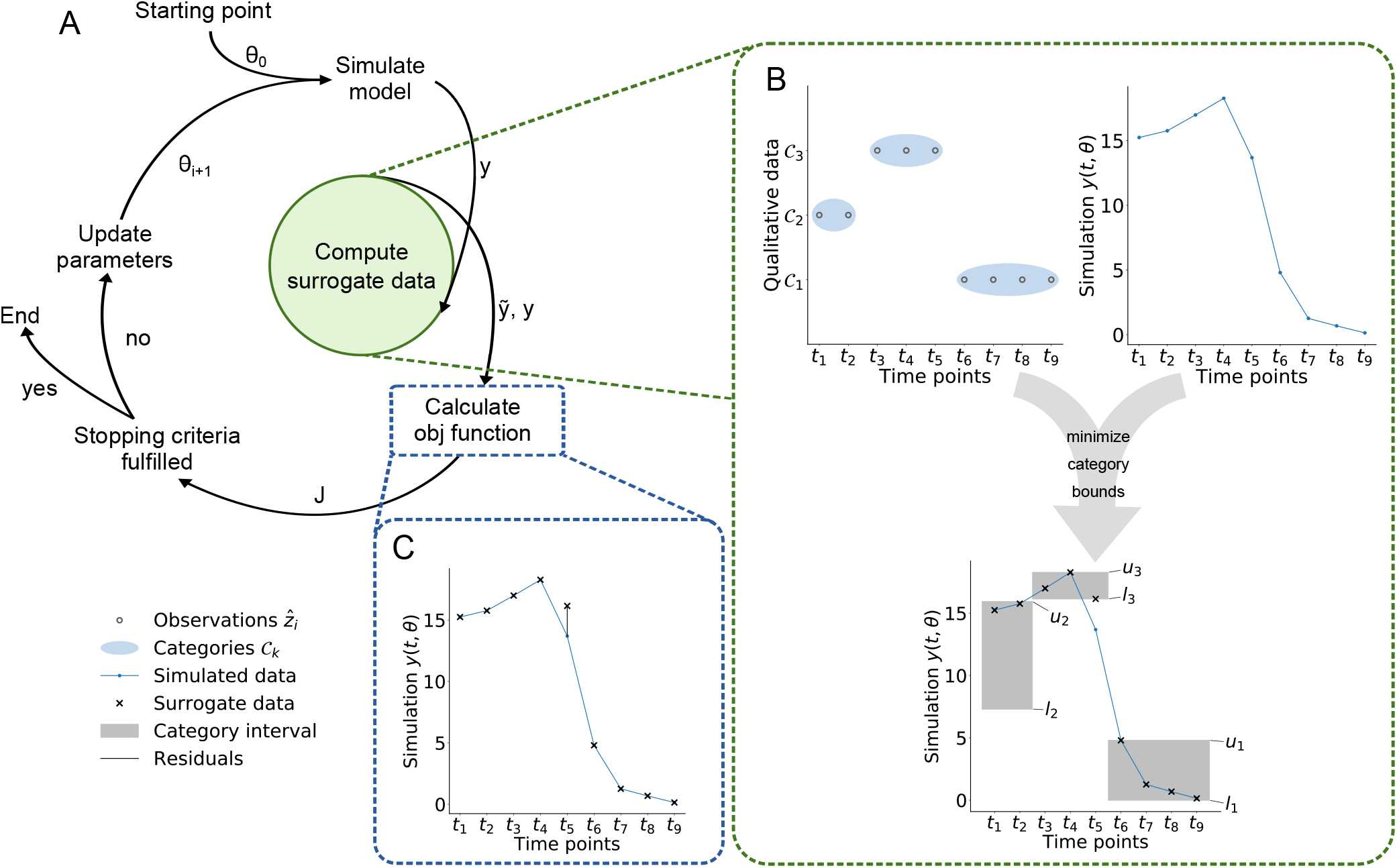
Illustration of the optimal scaling approach. (A) Individual steps of an optimization run. (B) Schematic of surrogate data calculation for a given simulation results *y*(*t,θ*) and set of qualitative data (with three categories). The interval between the optimized lower and upper bounds of the categories are indicated by grey areas. (C) Schematic of residuals used in the objective function for the parameter optimization.

Here, we build upon the optimal scaling method developed by Pargett & Umulis (2013) and Pargett *et al.* (2014) for dynamical systems. To accelerate the solution of the inner optimization, we first propose a reduced formulation which conserves the optimal points. Next, the reduced formulation is reparameterized to an unconstrained optimization problem, which can be solved more robustly as we demonstrate on three application examples. The approach is implemented in the Python Parameter EStimation TOolbox pyPESTO (Schälte *et al.*, 2019) and can be used with the parameter estimation data format PEtab (Weindl *et al.*, 2019) making it easily reusable.

## 2 Methods

### 2.1 Modeling of biochemical processes

We consider biochemical processes described by ordinary differential equations (ODEs) of the form:

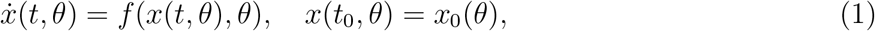

in which *x*(*t, θ*) ∈ ℝ^*n_x_*^ denotes the concentrations of biochemical species at time *t* and *f* : ℝ^*n_x_*^ × ℝ^*n_θ_*^ ↦ ℝ^*n_x_*^ the vector field describing their temporal evolution. The vector field is assumed to be Lipschitz continuous in *x* to ensure existence and uniqueness of the solutions. The vector *θ* ∈ ℝ^*n_θ_*^ comprises the unknown time-invariant parameters of the ODE (1). The function *x*_0_ : ℝ^*n_θ_*^ ↦ ℝ^*n_x_*^ provides the parameter-dependent initial condition at initial time *t*_0_, thereby allowing for steady state constraints (Rosenblatt *et al.*, 2016; Fiedler *et al.*, 2016).

### 2.2 Measurement process

We consider quantitative and qualitative measurement data. To allow for partial observations of the state x(t, θ), we define the observation function *h* : ℝ^*n_x_*^ × ℝ^*n_θ_*^ ↦ ℝ. The observable *y*(*t, θ*) ∈ ℝ is given by

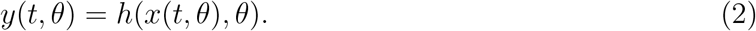

Examples for the observation function are *h*(*x, θ*) = *x*_1_ (absolute measurements of state variable *x*_1_), *h*(*x,θ*) = *x*_1_ + *x*_2_ (absolute measurements of the sum of state variables *x*_1_ and *x*_2_), and *h*(*x,θ*) = *θ*_1_*x*_1_ (relative measurements of state variable *x*_1_). Also saturation effects and more complex dependencies can be considered.

For ease of notation we consider in the main manuscript the case of a single observable and a single time-lapse experiment. The extension to multiple observables and multiple experiments (e.g. a dose-response curve) is straight forward.

**Quantitative data** are noise-corrupted observations of *y*(*t,θ*),

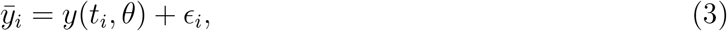

with time index *i* = 1,…, *N*. Here, we assume additive and normally distributed measurement noise 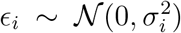 with standard deviation *σ_i_*. Alternatives are provided by Laplace and t-distributed measurement noise (Maier *et al.*, 2017).

**Qualitative data** are information about a readout *z*(*θ, t*) which is related to the observable *y*(*θ, t*). Yet, the mapping from the observable *y*(*θ, t*) to the measured quantity is not precisely known. For several experimental techniques only monotonicity of the mapping from *z*(*θ,t*) to *y*(*θ,t*) can be assumed. This means that an increase of the readout *z*(*θ,t*) implies an increase of the observable *y*(*θ,t*), but that *y*(*θ,t*) might increase without changing *z*(*θ,t*). This happens for instance if the readout is discrete or if there is a detection limit or detector saturation.

The measured (qualitative) readouts are potentially noise corrupted

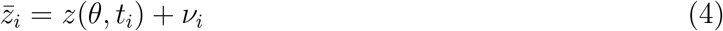

with measurement noise *ν_i_* and are either indistinguishable, i.e. 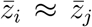, or ordered, 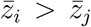 or 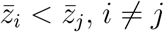. We follow the formulation by (Pargett & Umulis, 2013) and introduce categories 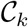, *k* = 1,…, *K*, which are without loss of generality assumed to be ordered as 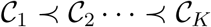. The categories contain observations, which are indistinguishable from each other, i.e. 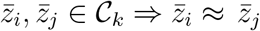. Observations from different categories can be distinguished by the ordering of the categories. The index of the category to which observation 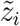 belongs is denoted by *k*(*i*). An illustration of qualitative data is shown in Figure 1B.

### 2.3 Parameter estimation

The unknown parameters *θ* of the ODE model (1) have to be inferred from the available quantitative and qualitative data.

For **quantitative data**, parameter estimates are usually computed by minimizing the difference between the data and the model simulation. The difference is commonly formulated in terms of the negative log-likelihood or the negative log-posterior functions (Raue *et al.*, 2013b). Here, we considered normally distributed measurement noise with known standard deviation. In this case the negative log-likelihood function is a weighted least squares objective function. The corresponding optimization problem is

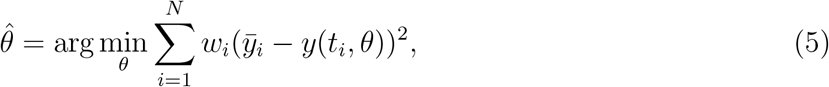

with quantitative data 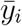, model simulation *y*(*t_i_,θ*) and weights 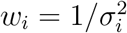. Multi-start local optimization has been shown to be a competitive method for solving these types of ODE-constrained optimization problems (Raue *et al.*, 2013a; Villaverde *et al.*, 2018).

For **qualitative data**, parameter estimates can be computed using the optimal scaling approach (Pargett *et al.*, 2014) (Figure 1A-C). This approach addresses the problem that the mapping from quantitative simulation to qualitative data is unknown by introducing quantitative surrogate data 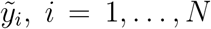. These surrogate data provide the best agreement with the model simulation within the constraints provided by the qualitative data (Figure 1B), i.e. the information about the category of a data point and its (qualitative) relation to other data points. For a given parameter *θ* and corresponding model simulation *y*(*t, θ*), the surrogate data are obtained by solving the optimization problem

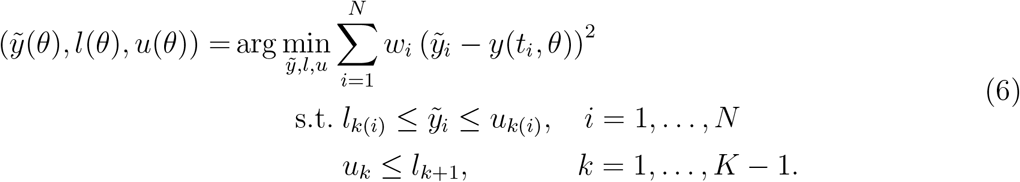

The qualitative information is enforced by restricting the surrogate data for observations in category 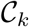 to the interval [*l_k_*, *u_k_*], with lower bound *l_k_* ∈ ℝ and upper bound *u_k_* ∈ ℝ. To ensure that categories are distinguishable, the upper bound of category 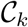 has to be lower than the lower bound of category 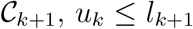. The category for the *i*-th observation is encoded in the index mapping *k*(*i*). The weights in the objective function are set to

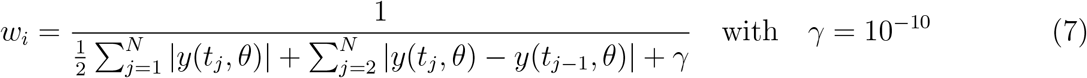

which is similar to the choice by Pargett *et al.* (2014). This choice of the weight ensures that the values of the observable do not tend to zero, thereby improving numerical stability compared to a choice which is independent of the simulation. The second summand penalizes flat simulations and *γ* is chosen such that *w* is still evaluable for simulations equal to zero.

To estimate the parameters *θ*, the distance between model simulation and optimal surrogate data (Figure 1C) is minimized

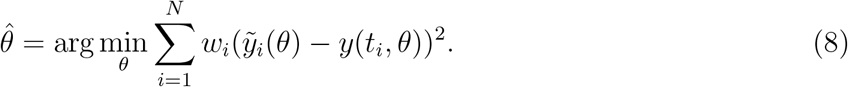

As the surrogate data possess for all *θ* the correct qualitative characteristics, the minimization of the objective function 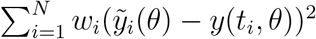 yields a sequence of points which approaches the measured qualitative dynamics. If the model simulations show the correct qualitative behaviour, the objective function becomes zero.

The surrogate data depend on the parameter-dependent model simulation *y*(*t_i_,θ*). Therefore, the optimization of the surrogate data is nested in the optimization of the parameters *θ* and has to be performed in each optimization step. To accelerate this process, Pargett *et al.* (2014) employed that the optimal surrogate data can be computed from the optimal category bounds *u*(*θ*) and *l*(*θ*):

(Case 1) If the model simulation *y*(*t_i_,θ*) is smaller than the lower bound *l*_*k*(*i*)_(*θ*), the surrogate data are set to the smallest feasible value to minimize the difference, i.e. 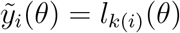.
(Case 2) If the model simulation *y*(*t_i_, θ*) is larger than the upper bound *u*_*k*(*i*)_(*θ*), the surrogate data are set to the largest feasible value to minimize the difference, i.e. 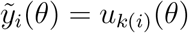.
(Case 3) If the model simulation *y*(*t_i_,θ*) is in the interval [*l*_*k*(*i*)_(*θ*), *u*_*k*(*i*)_(*θ*)], then the surrogate data are set to 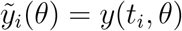. In this case the error is zero.

These analytical results provide a construction rule for the surrogate data:

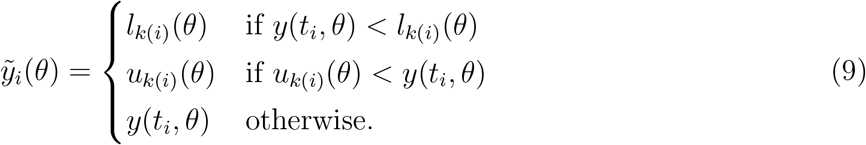

Using this construction rule, the category bounds can be computed using the optimization problem:

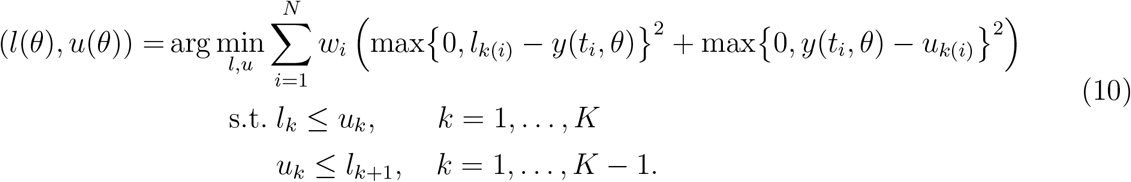

In the considered objective function, the term max{0, *l*_*k*(*i*)_ – *y*(*t_i_, θ*)}^2^ vanishes in Case 2 and 3 while the term max{0, *y*(*t_i_, θ*) – *u*_*k*(*i*)_}^2^ vanishes in Case 1 and 3. Accordingly, the objective function uses the analytical results for the optimal surrogate data and its minimizer provides the optimal lower and upper bounds for the categories. Hence, in the optimal scaling approach (Figure 1A), solving (6) can be replaced by solving (10) and evaluating the optimal surrogate data using (9).

In the optimization problems (6) and (10), the qualitative data provide only limited information about the lower bound *l*_1_ of category 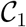 and the upper bound *u_K_* of category 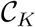. The lower bound *l*_1_ may be set to any value smaller or equal to the minimum of *y*(*t_i_,θ*), *l*_1_ ≤ min_*i*_ *y*(*t_i_,θ*), and the upper bound *u_K_* may be set to any value greater or equal to the maximum of *y*(*t_i_,θ*), *u_K_* ≥ max_*i*_ *y*(*t_i_,θ*).

### 2.4 Acceleration of surrogate data calculation

The surrogate data calculation proposed by Pargett *et al.* (2014) reduces the number of optimization variables from *N* + 2(*K* – 1) to 2(*K* – 1). Yet, the calculation of the surrogate data is for many application problems still the most time-consuming process within the parameter estimation. Here, we propose two reformulations to accelerate the surrogate data calculation.

The first reformulation is based on our empirical observation that the gaps between lower and upper bounds of adjacent categories are often estimated as small as possible. Our analysis of the phenomenon revealed:

#### Lemma 1.

*The optimization problem* (10) *possesses an optimal solution* (*l*\*,*u*\*) *with* 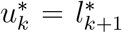 *for k* = 1, …, *K* – 1.

*Proof.* Assume there is an optimal solution (*l′, u′*) with a non-zero gap between adjacent lower and upper bounds. Without loss of generality, we assume that 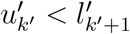.

For all observations *i* ∈ {1,…, *N*} with *k*(*i*) = *k*′ it has to hold that 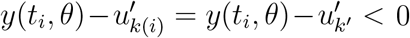. Otherwise, the objective function could be decreased by setting 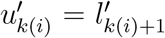 as the objective function summand 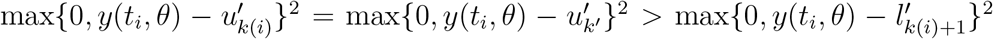 would decrease. This would imply that (*l′*, *u′*) is not an optimal solution.

As 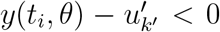 the corresponding objective function summands are zero, 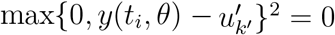. This does not change if 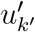, is increased to 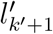.

This proof also holds for other weighting functions *w_i_* (*y*, *l′*, *u′*) as well as constant weights *w_i_*.

Lemma 1 implies that among the optimal solutions of (10), there is at least one for which the lower and upper bounds of adjacent categories are identical (see also Figure 2). Accordingly, an optimal solution of (10) can be computed by solving a reduced problem:

**Figure 2:**
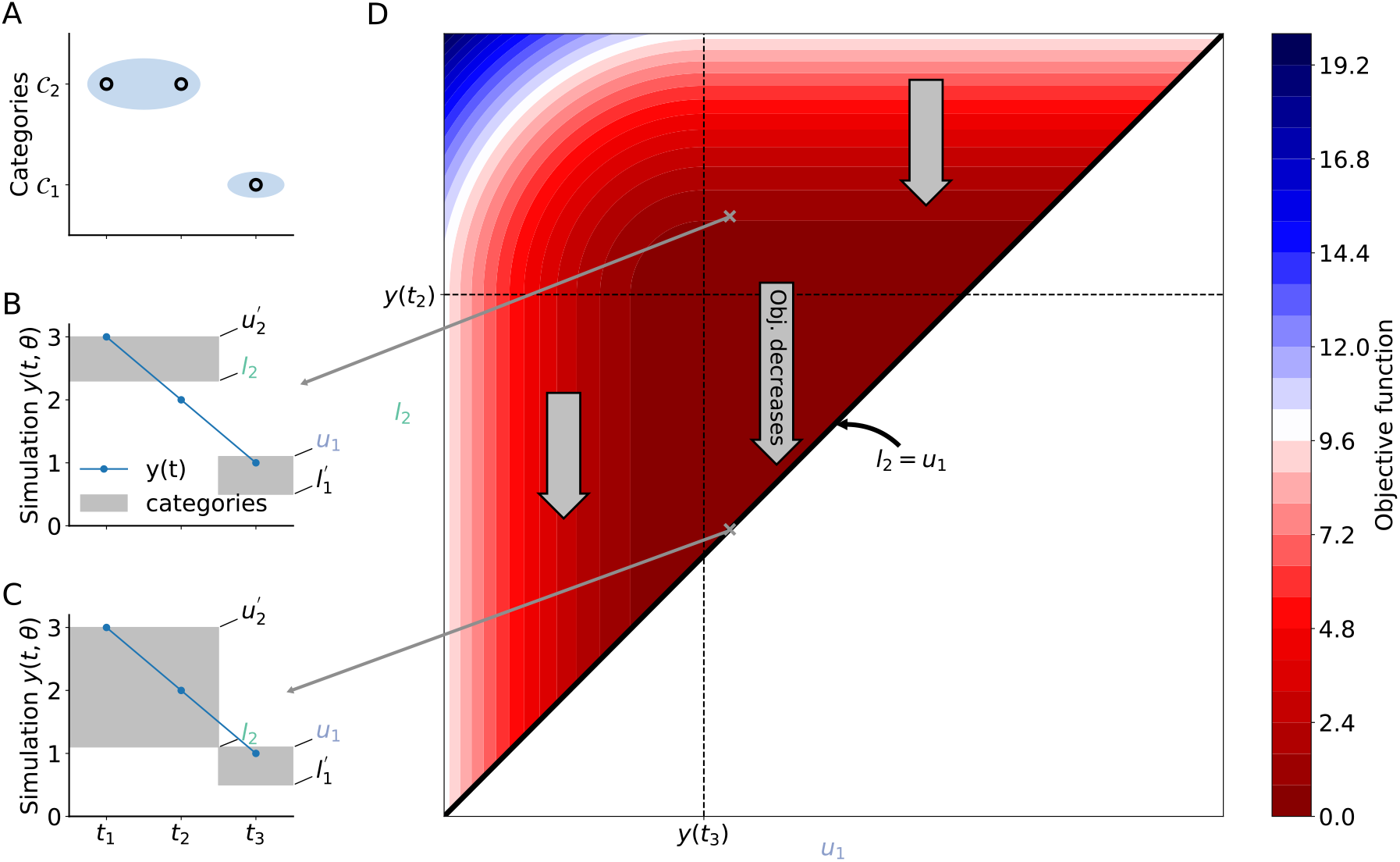
Schematic representation of the reduction of the inner optimization problem. (A) Example of qualitative data with two categories and three observations. (B & C) Simulated data with category bounds shown in gray for two different category bounds. (D) Objective function landscape for upper bound of 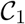 and lower bound of 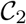, showing that the objective function decreases (or stays constant), when decreasing the gap between the two category intervals. By setting *l*_2_ = *u*_1_ the minimal objective function is achieved.

#### Theorem 2.

*An optimal solution of the optimization problem* (10) *is obtained by solving*

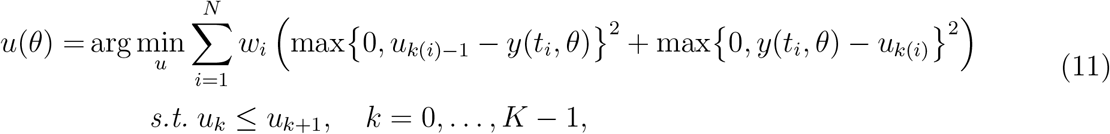

*for u*_0_ = min_*i*_ *y*(*t_i_, θ*) *and u_K_* = max_*i*_ *y*(*t_i_, θ*), *and setting l*_*k*+1_(*θ*) = *u_k_* (*θ*) *for k* = 1,…,*K* – 1.

*Proof.* Optimization problem (11) is obtained by substituting *l*_*k*+1_ with *u_k_* in optimization problem (10) and removing trivially fulfilled constraints. The reduced optimization problem obtained by the substitution effectively solves (10) on the subspace *l*_*k*+1_ = *u_k_*, which contains one of the optimal solutions (Lemma 1).

We note that *u*_0_ is an auxiliary variable used to simplify the notation, not the bound of an additional category.

The reduced optimization problem (11) possesses *K* – 1 optimization variables. Hence, the number of optimization variables is reduced by a factor of two compared to the available formulation (11). This should accelerate the optimization. Yet, as the objective function is nonlinear and as we have linear inequality constraints, the availability of optimization methods is limited.

The second reformulation is based on our empirical finding that available solvers for nonlinear optimization problems with box constrained optimization variables are often computationally more efficient than those for general linear inequality constraints. To this end, we introduce the vector of differences between the upper bounds of adjacent categories, *d_k_* ≔ *u_k_* – *u*_*k*−1_. Using this difference, the category bounds can be written as 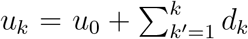. The auxiliary variable *u*_0_ can be set to some value lower or equal to the minimum of *y*(*θ,t_i_*), e.g. *u*_0_ = min_*i*_ *y*(*θ,t_i_*). The reformulation of the reduced optimization problem using the differences yields

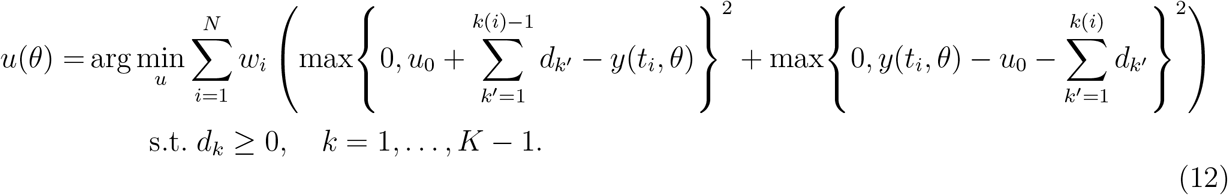

This optimization problem contains only positivity constraints for the optimization variables. Hence, a broader spectrum of nonlinear optimization algorithms can be employed.

To select appropriate numerical optimization algorithms, we analyzed the properties of the optimization problems. We found that:

#### Theorem 3.

*The optimization problems* (10), (11), *and* (12) *are convex*.

*Proof.* The objective functions of the respective optimization problems are sums of convex functions of the lower bounds *l*, the upper bounds *u* and/or the differences *d*. As the sum of convex functions is itself convex (Boyd & Vandenberghe, 2004, Section 3.2), the overall objective function is convex. In combination with linear inequality constraints, this implies that the optimization problem is convex.

Convex optimization problems only possess one optimum. Hence, local optimization methods should – in theory – converge to the optimal solution.

### 2.5 Category and gap sizes

To ensure that qualitatively different readouts are related to non-negligible quantitative differences, Pargett *et al.* (2014) enforced a minimal size *s* ∈ ℝ_+_ for each category and a minimal gap *g* ∈ ℝ_+_ between categories. Therefore, the constraints were modified, yielding the optimization problem

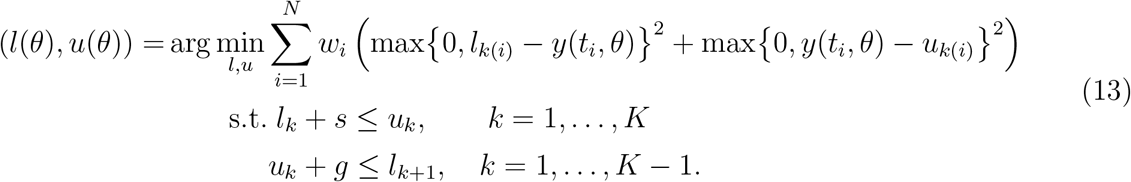

For this optimization problem it can be shown that the optimal values for lower and upper bounds are in the interval [min_*i*_ *y*(*t_i_,θ*) – *K*(*g* + *s*), max_*i*_ *y*(*t_i_,θ*) + *K*(*g* + *s*)]. Outside of the interval the objective function is – independent of the specific simulation results – increasing. Accordingly, one can set *l*_1_ = min_*i*_ *y*(*t_i_,θ*) – *K*(*g* + *s*) and *u_K_* = max_*i*_ *y*(*t_i_,θ*) + *K*(*g* + *s*).

For optimization problem (13) it can be shown that there exists an optimal solution with *l*_*k*+1_ = *u_k_* + *g*. This is a straight extension of Lemma 1 and provides the basis for reformulation of (13) similar to results presented in Section 2.4:

#### Theorem 4.

*An optimal solution of the optimization problem* (13) *is obtained by solving*

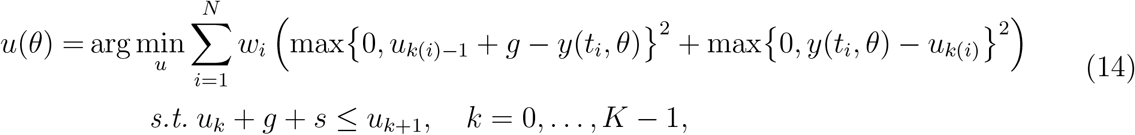

*for u*_0_ = min_*i*_ *y*(*t_i_, θ*) – *K*(*g* + *s*) *and u_K_* = max_*i*_ *y*(*t_i_, θ*) + *K*(*g* + *s*), *and setting l*_*k*+1_(*θ*) = *u_k_*(*θ*) + *g for k* = 1, …, *K* – 1.

The proof of Theorem 4 is analogue to the proof of Theorem 2, which is a special case of this result for *s* = *g* = 0.

The reduced optimization problem (14) can be again reformulated to replace the linear inequality constraints with positivity constraints. Here, we use the difference between uk and its minimal value given *u*_*k*−1_ and the required gaps, *d_k_* ≔ *u_k_* – (*u*_*k*−1_ + *g* + *s*), as new optimization variables. This yields

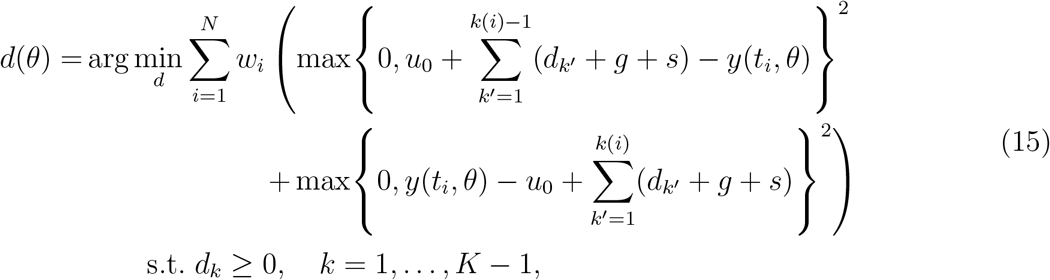

with some *u*_0_ ≤ min_*i*_ *y*(*t_i_,θ*) – *K*(*g* + *s*).

The optimization problems (13), (14) and (15) with constraints on category and gap sizes are also convex. To show this the proof of Theorem 3 can be reused.

#### Remark 5.

*In Section 2.3-2.5, we use the structure of the optimization problems to provide conservative bounds for u*_0_, *u_K_ and l*_1_. *In practice, these bounds might be tightened using additional information, e.g., that the data are non-negative.*

## 3 Application

To evaluate the optimal scaling approach with the reformulated surrogate data calculation, we implemented the approach and compared accuracy and computation time to those of available methods.

### 3.1 Implementation

We implemented the optimal scaling approach for parameter estimation with qualitative data in pyPESTO (Schälte *et al.*, 2019). Our implementation allows to choose between surrogate data calculation using

- the **standard** optimization problem (13),
- the **reduced** optimization problem (14), and
- the **reparameterized** reduced optimization problem (15)

for the calculation of the category bounds.

For the surrogate data calculation we employed two optimization algorithms: For the standard and the reduced optimization problems with linear inequality constraints we used the Sequential Least Squares Programming (SLSQP) algorithm. For the reparameterized reduced optimization problem with box constraints we used the L-BFGS-B algorithm. These optimization algorithms are implemented in the Python package SciPy (Jones *et al.*, 2001). We allowed for a maximum of 2000 iterations and set the function tolerance to 10^−10^. For the selection of the minimal gaps between categories and minimal category sizes we follow the recommendation of Pargett *et al.* (2014) but additionally enforce a minimum of *ϵ* = 10^−16^:

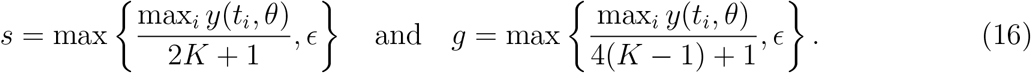

The minimum value *ϵ* facilitates the mitigation of numerical integration errors for the ODE model. Initial guesses of the bounds are placed between 0 and max_*i*_ *y*(*t_i_,θ*) + *s*, and reparameterized to obtain the starting points for the reparameterized reduced formulation. If the calculation of the category bounds fails, the objective function value of the outer loop is set to NaN.

For the calculation of the parameters *θ* using the optimization problem (8), we employed the Nelder-Mead and Powell algorithm. These gradient-free algorithms are interfaced through pyPESTO and turned out the be more reliable than the available gradient-based methods. The reason was probably that for the specific problem structure, finite difference approximations of the gradient were inaccurate and sensitivity-based gradient calculation is not implemented. As stopping criteria for the outer optimization, we used an absolute function tolerance of 10^−10^ and a maximum of 500 number of iterations and function evaluations. The optimization was performed in log-space.

For the numerical simulation of the ODE models we used the Advanced Multilanguage Interface to CVODES and IDAS (AMICI) (Fröhlich *et al.*, 2017), which internally exploits the Sundials solver package (Hindmarsh *et al.*, 2005). We set the absolute tolerance to 10^−6^ and the relative tolerance to 10^−8^.

The source code of all performed analysis will be made available upon final publication. The implementation of the optimal scaling approaches is available at https://github.com/ICB-DCM/pyPESTO/tree/feature_ordinal.

### 3.2 Test problems

For the evaluation of the proposed methods, we considered three published models. These models possess 5 to 14 state variables, 2 to 18 unknown parameters, and 1 to 8 observables. An overview about the model properties is provided in Table 1.

**Table 1:** Overview over the considered models and their properties, as well as the corresponding datasets.

| Model | RAF<br>inhibition | STAT5<br>dimerization | IL13-induced<br>signaling |
| --- | --- | --- | --- |
| number of state variables, $n_x$ | 5 | 8 | 14 |
| number of parameters, $n_\theta$ | 2 | 6 | 18 |
| number of observables | 1 | 3 | 8 |
| number of data points, $N$ | 9 | 48 | 205 |
| number of categories, $K$ | 2–9 | $3 \times 16$ | 6–38 |
| Reference | Mitra *et al.*<br>(2018) | Boehm *et al.*<br>(2014) | Raia *et al.*<br>(2011) |

The model of **RAF inhibition** used by Mitra *et al.* (2018) is used as an illustration example. It comprises two unknown parameters and we consider 9 simulated data points, discretized in 2 to 9 categories.

The **STAT5 dimerization** model by Boehm *et al.* (2014) is considered as a small application problem. This model describes the homo- and heterodimerization of the transcription factor isoforms STAT5A and STAT5B using 6 unknown parameters. It has 3 observables for each of which 16 quantitative measurements are available. For the evaluation of the proposed optimal scaling approach we consider as qualitative data the ordering of the measured values. As the values of different observables is not necessarily comparable, separate orderings are used for the observables, yielding 3 × 16 categories, and the surrogate data calculation is performed separately for each observable.

The model of **IL13-induced signaling** by Raia *et al.* (2011) is considered as a larger application example. This model describes IL13-induced signaling in Hodgkin and Primary Mediastinal B-Cell Lymphoma. It comprises 18 unknown parameters and 7 observables, for which 6–38 quantitative measurements are available. As qualitative data we consider again the ordering of the measured values.

In this study, we considered application examples for which quantitative measurements are available and which are included in a collection of benchmark problems for parameter estimation, which facilitates easy reusability (Hass *et al.*, 2019). This enables a comparison of parameter estimation using quantitative and qualitative data. For more details on the models we refer to the original publications.

### 3.3 Convexity, optimality and scalability

To verify the theoretical finding that optimization problems for the calculation of the category bounds are convex, we performed multi-start local optimization for the model of RAF inhibition (Figure 3A). The waterfall plot reveals that for this model all starts converged to the same objective function value (Figure 3B). This is in line with our our theoretical findings.

**Figure 3:**
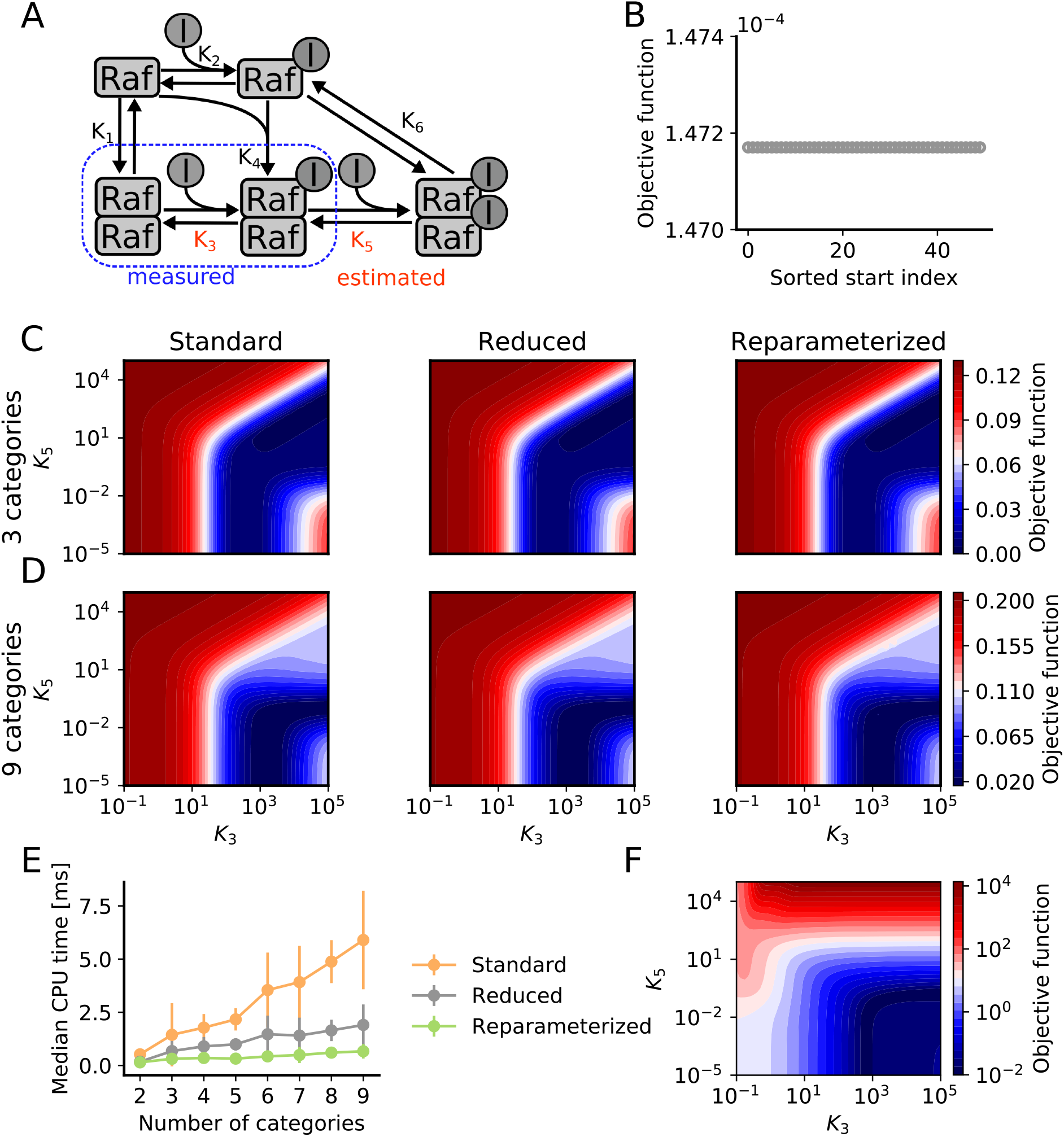
Comparison of standard and reduced formulations for the calculation of the surrogate data for the model of RAF inhibition. (A) Illustration of the model. (B) Waterfall plot of multi-start local optimization results for surrogate data calculation with 3 categories for the reduced formulation. The objective function was evaluated at the model parameters *K*_3_ = 4000 and *K*_5_ = 0.1. (C,D) Objective function landscape for qualitative data with (C) 3 categories and (D) 9 categories. (E) Computation time for the calculation of the surrogate data. (F) Objective function landscape for quantitative data.

To confirm that the reduced formulations provide optimal surrogate data, we evaluated the objective function using the standard and the reduced optimization problems. Since this model has only two unknown parameters, we studied the complete objective function landscape for the dataset with 3 and 9 categories (Figure 3C,D). The numerical results confirm that the objective function values obtained with the different approaches are identical.

Despite the convexity of the optimization problems, the computational complexity substantially increases with the number of categories (Figure 3E). While the absolute computation time of reduced and reparameterized reduced formulation is lower than for the standard formulation, the scaling behaviour is comparable. The computation time depends linearly on the number of categories.

### 3.4 Information content

Qualitative data are often assumed to provide a limited amount of information. To assess this hypothesis, we studied the objective function for the model of RAF inhibition for qualitative data with different numbers of categories (Figure 3C,D) as well as quantitative data (Figure 3F). Interestingly, the objective function landscape for qualitative data hardly depends on the number of categories and closely resembles the objective function landscape for quantitative data. This implies that qualitative data can be almost as informative as quantitative data. This is corroborated by the objective function profiles we computed (Figure 4), which can be used for uncertainty analysis (Raue *et al.*, 2009).

**Figure 4:**
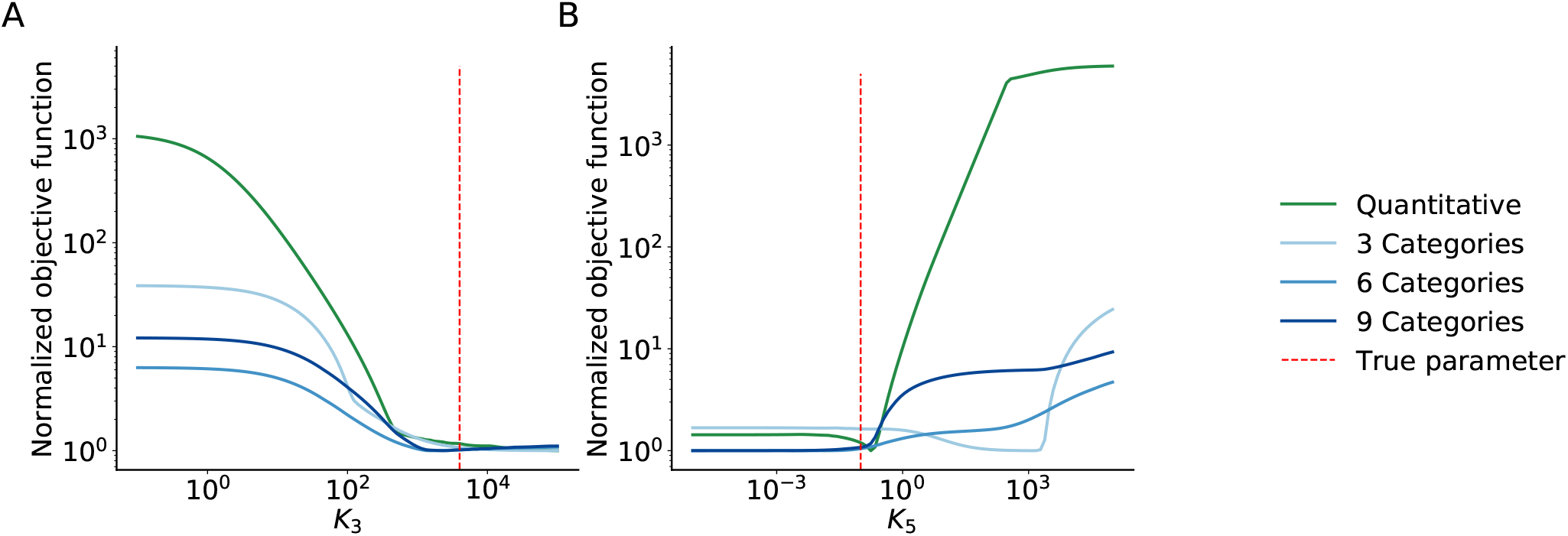
Objective function profiles for the model of RAF inhibition. (A & B) Profiles for qualitative data with 3, 6 and 9 categories and quantitative data for Parameter *K*_3_ (A) and *K*_5_ (B).

### 3.5 Robustness and efficiency

The reduced formulations for the surrogate data calculation possess only half as many optimization variables as the standard formulation, and the reparameterized reduced formulation possesses only positivity constraints. To evaluate the practical impact of these reformulations, we solved the respective optimization problems for the models of STAT5 dimerization and IL13-induced signaling. For each model, 150 parameter vectors were sampled and the corresponding category bounds were computed.

Although the considered inner optimization problems are convex, the considered optimization algorithms provided different results for the different formulations (Figure 5A). To our surprise, numerical optimization often failed to provide appropriate category bounds when using the standard formulation (Figure 5B). For the model of IL13-induced signaling, only 37% of the optimizations with the standard and 36% with the reduced formulation were successful. For the remaining ones, the optimizer failed for different reasons. This problem was not observed for the reparameterized reduced formulation, probably because the optimization algorithm we can employ for this problem is more reliable.

**Figure 5:**
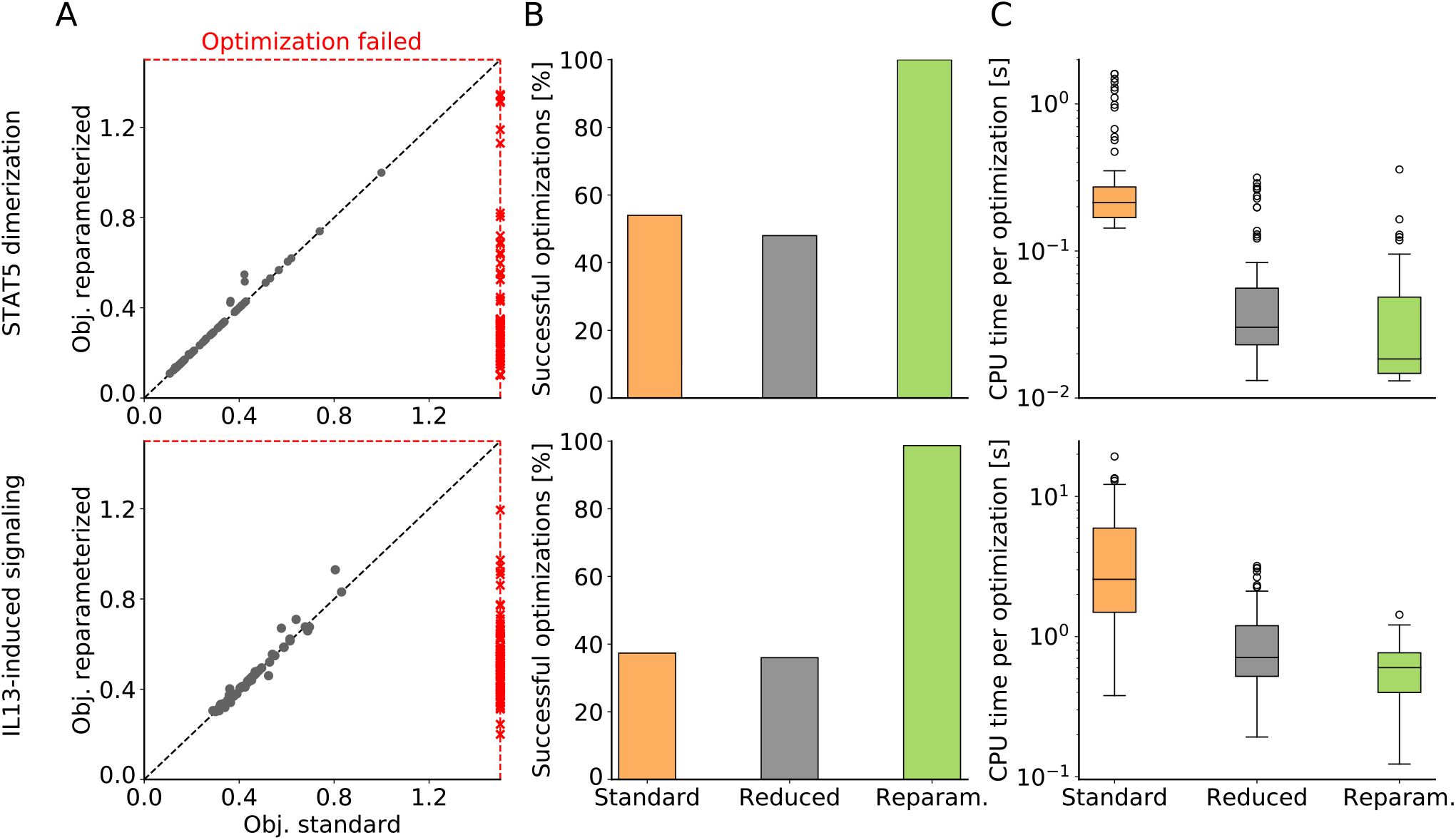
Computation time and robustness of standard and reduced formulations for the calculation of the surrogate data. (A) Scatter plot with final objective function values obtained using standard and reparameterized reduced formulation for 150 randomly sampled parameter values. Black dots correspond to starting points for optimization for which standard and reparameterized reduced formulation was successful, while red crosses indicate that the corresponding optimization failed. (B) Percentage of successful calculations of the surrogate data. (C) Computation times for standard, reduced and reparameterized reduced formulation. Only computation times for successful evaluations are shown.

For the sampled parameter vectors for which numerical optimization for all formulations was successful, the computation time for the reduced and the reparameterized reduced formulation is substantially lower than for the standard formulation (Figure 5C). We observed median and mean speedups of 11.5 and 18.9 respectively for the model of STAT dimerization and 4.2 and 7.4 for the model of IL13-induced signaling for the reparameterized reduced formulation compared to the standard formulation. Hence, the proposed formulations allow for more robust and more efficient calculation of the surrogate data.

### 3.6 Overall performance

The calculation of the category bounds and the surrogate data is only one step in the parameter estimation loop (Figure 1). To assess the overall performance of parameter optimization using standard and reduced formulations, we performed a multi-start local optimization using gradient-free optimizers Nelder-Mead and Powell.

The results of the multi-start local optimization reveal that standard and reduced formulations yield similar final objective function values (Figure 6A). In all cases except for the model of STAT5 dimerization with Nelder-Mead algorithm, the reparameterized formulation achieved slightly better objective function values. This might be due to the improved robustness of the evaluation of the inner problem demonstrated in Section 3.5.

**Figure 6:**
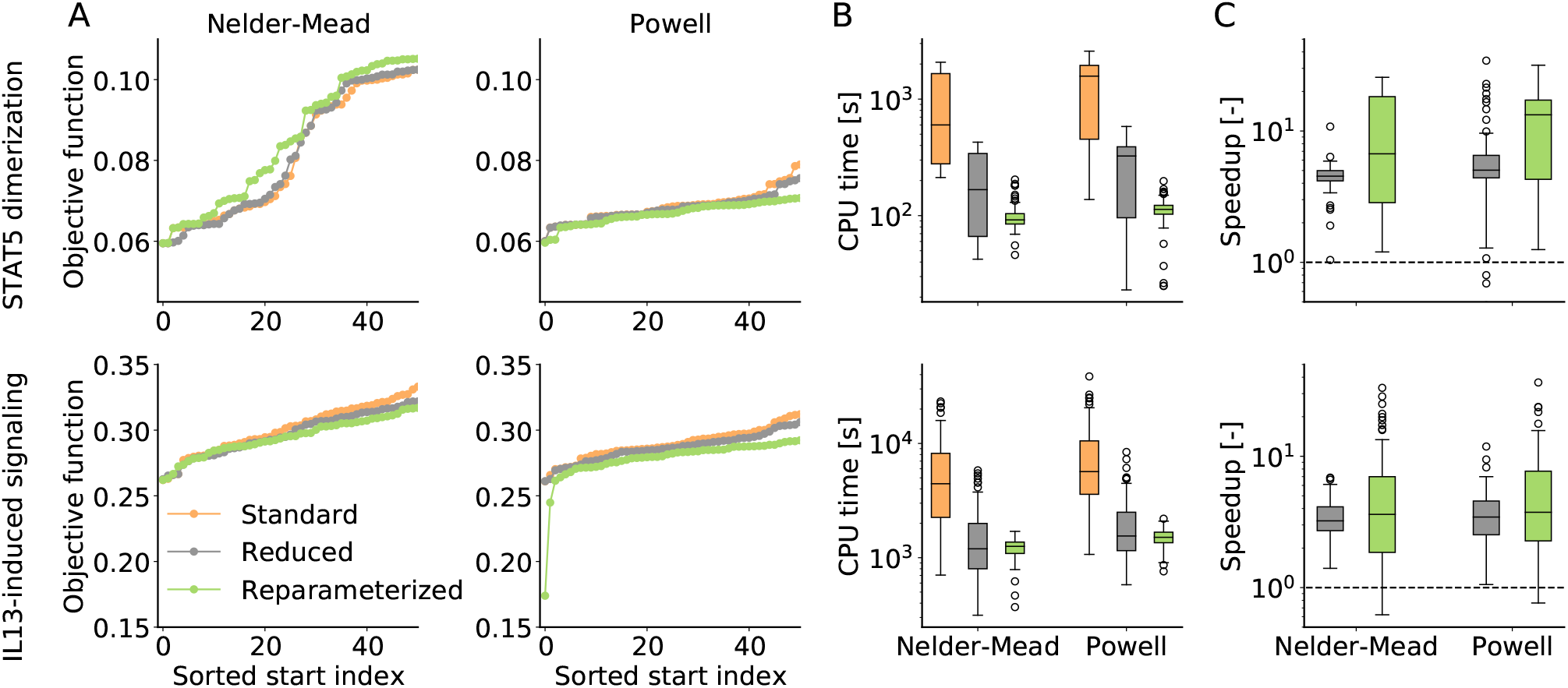
Parameter optimization for the models of STAT5 dimerization and IL13-induced signaling. (A) Waterfall plots for different combinations of model, optimization algorithm and formulation of the surrogate data calculations. The best 50 starts out of total of about 100 runs are shown. (B) Computation times for the different combinations. (C) Speedups achieved using the reduced formulations. Above the dashed line the use of the reformulation was computationally more efficient and below the use of the standard formulation.

As the calculation of the surrogate data requires a substantial amount of the overall computation time, the improved efficiency of the reduced formulations demonstrated in Section 3.5 decreases the computation time (Figure 6B). On average we observe a 5–10-fold reduction of the computation times for a local optimization (Figure 6C).

## 4 Discussion

Measurements that provide qualitative information are common in biology. Yet, only few approaches exist to incorporate qualitative measurements in the development of dynamic models (Mitra *et al.*, 2018; Pargett *et al.*, 2014) and these approaches are computationally demanding. Here, we built upon the optimal scaling approach introduced in Pargett *et al.* (2014) and show that this approach can be reformulated to a problem with a reduced number of optimization variables.

We evaluated the proposed reparameterized formulation of the optimal scaling approach using three application examples and observed a 3- to 10-fold speedup. The speedup increased with the size of the dataset per observable. Even more important than the speedup could be the finding that proposed optimal scaling approach is more robust and yields often better final objective function values. These benefits were independent of the optimization algorithm.

Open questions for the proposed approach include the choice of the weighting factors *w_i_*, the minimal gap between categories *g* and the minimal size of categories *s*. We observed that for the latter to the suggestions found in the literature are often not ideal. Furthermore, in this study, we only used gradient-free optimization algorithms, although optimization algorithms exploiting gradient information often proved to be more efficient and reliable (Raue *et al.*, 2013a; Villaverde *et al.*, 2018; Schälte *et al.*, 2018). To further improve the parameter estimation, the gradient of the objective function could be employed, which requires the sensitivity of the parameter-dependent surrogate data. As the surrogate data are the solution to the optimization problem (6), their sensitivity is the sensitivity of this optimal solution with respect to the parameters. This sensitivity can be determined by differentiating the Karush-Kuhn-Tucker condition for (6) with respect to the parameters and solving it for the gradients. The respective equations are provided in Appendix A.1, but an evaluation needs to be performed.

As qualitative data provide less information about the dynamical system than quantitative measurements, identifiability is a key concern. Unfortunately, established methods and tools for structural identifiability analysis (Chis *et al.*, 2011; Ligon *et al.*, 2018) are not applicable to the problem class. Furthermore, while the optimal scaling approach can be easily used for profile calculation (Raue *et al.*, 2009) (see our results for the model of the RAF inhibition), the statistical interpretation of objective function differences is unclear. A first Bayesian formulation has been proposed (Mitra *et al.*, 2019), but the statistical interpretation is not completely clear. A proper statistical formulation would also benefit the integration of qualitative and quantitative data (Mitra *et al.*, 2018), and might improve parameter identifiability.

We conclude that the ability to use qualitative information is very important, but that there are many open problems. We provide an improved optimal scaling approach for dynamical systems and a corresponding open-source implementation. We expect that this will contribute to the further development of methods for the analysis of qualitative data.

## A Appendix

### A.1 Gradient of optimal surrogate data

To facilitate the use of gradient-based optimization algorithms for the parameter optimization problem with qualitative data (8), the gradient of the surrogate data with respect to the parameter vector θ is required. This is the gradient of the optimal solution of the optimization problem (6), which can be written as:

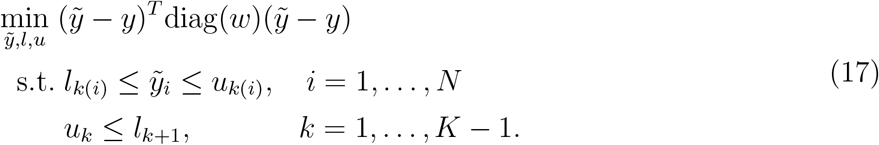

or more generally as

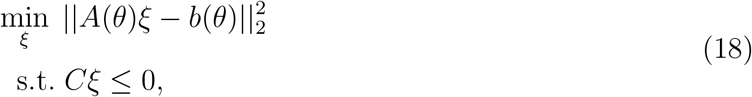

with 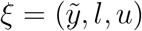.

The Lagrange function for (18) is

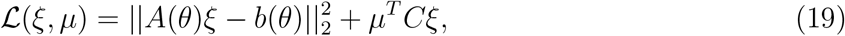

with the Lagrangian multiplier *μ* ≥ 0. This yields the first order optimality condition

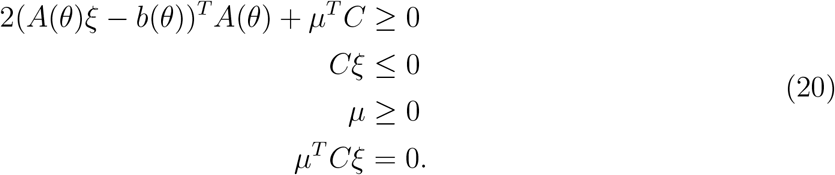

This system of equations describes the dependence of the optimal surrogate data and category bound (collected in *ξ*) as well as the Lagrange multiplier on *θ*. Therefore, we use the notation *ξ* ≔ *ξ*(*θ*) and *μ* ≔ *μ*(*θ*). The differentiation of (20) with respect to *θ* yields the derivative of the optimal surrogate data:

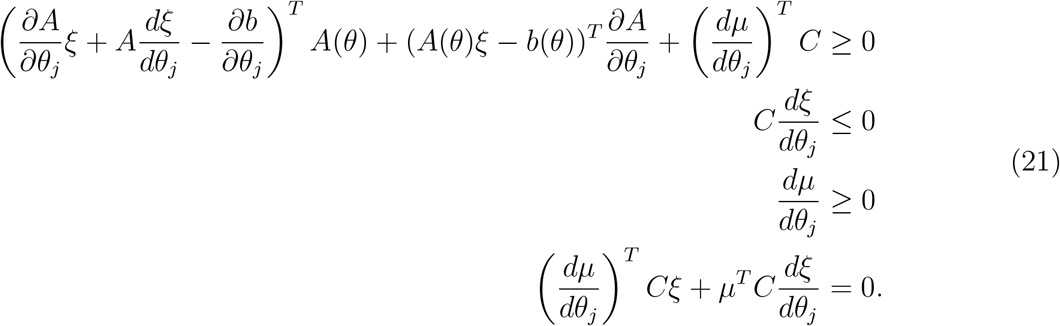

As *ξ* and *μ* are available, the system can be solved for 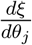 and 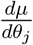 for *j* = 1,…, to obtain the derivatives. As the system is linear, efficient numerical methods are available.

## Author Contributions

L.S., J.H. derived the theoretical foundation; L.S. wrote the implementations and performed the case study; L.S., J.H. and D.W. analyzed the results; All authors wrote and approved the final manuscript.

